# *TranSyT*, an innovative framework for identifying transport systems

**DOI:** 10.1101/2021.04.29.441738

**Authors:** Davide Lagoa, José P. Faria, Filipe Liu, Emanuel Cunha, Christopher S. Henry, Oscar Dias

## Abstract

The importance and rate of development of genome-scale metabolic models have been growing for the last years, increasing the demand for software solutions that automate several steps of this process. However, since TRIAGE’s release, software development for automatic integration of transport reactions into models has stalled. Here we present the Transport Systems Tracker (*TranSyT*), the next iteration of TRIAGE. Unlike its predecessor, *TranSyT* does not rely on manual curation to expand its internal database, derived from highly-curated records retrieved from the Transporters Classification Database and complemented with information from other data sources. *TranSyT* compiles information regarding transporters families, transport proteins, and derives reactions into its internal database, making it available for rapid annotation of complete genomes. All transport reactions have GPR associations and can be exported with identifiers from four different metabolite databases. *TranSyT* is currently available as a plugin for *merlin* v4.0 and an app for KBase.

## INTRODUCTION

In the last decades, the number of sequenced genomes has grown exponentially. Several publicly available Bioinformatics tools for reconstructing high-quality Genome-Scale Metabolic (GSM) models were developed or improved to keep up with this progress. However, there is limited availability of reliable tools for automatic annotation of transporter systems (1). Databases like the Transporter Classification Database (TCDB) (2), TransportDB 2.0 (3), ARAMEMNON (4), YTPdb (5), and ABCdb (6) contain data regarding transport proteins. TCDB was already used as the main source of information by the Transport Reactions Annotation and Generation (*TRIAGE*) (7), the predecessor of *TranSyT* because of its importance and curated content. TCDB contains structural, functional, mechanistic, evolutionary, and medical information about transport systems. Their authors also proposed the Transport Classification (TC) system, the only classification adopted by the International Union of Biochemistry and Molecular Biology (IUBMB) for transporters until August 2018, when the Enzyme Commission (EC) number 7, Translocases, was added by the organisation.

There have been several approaches in the development of software to tackle the different problems of transporters annotation. Regarding the automatic assembly of transport reactions, Lee et al. (8) developed the Transport Inference Parser (TIP) in 2008, a method for building transport reactions based on an organism’s genome annotation through textual analysis techniques. It analyses the name and function of each protein, thus inferring the transport reaction promoted by them.

Regarding the integration of transport information in GSM models, tools like ModelSEED (9) and Pathway Tools (10), predict transporters based on the Rapid Annotation using Subsystem Technology (RAST) (11) functional annotations to develop models and add spontaneous reactions to fill in pathways when necessary. Other tools, such as Reconstruction, Analysis and Visualization of Metabolic Networks (RAVEN) (12) or SuBliMinaL (13), generate transport reactions based on information retrieved from databases. RAVEN automatically retrieves data from the Kyoto Encyclopedia of Genes and Genomes (KEGG) (14) and the Braunschweig Enzyme Database (BRENDA) (15) or existing models of closely related organisms, based on the assumption that related organisms share metabolic capabilities. Additionally, RAVEN allows performing manual integration of transport reactions and filling gaps using KEGG Orthology (KO) identifiers to guarantee a functional network. SuBliMinaL (13) optionally provides a default set of transporters, retrieved from the Biochemical, Genetic and Genomic (BiGG) (16) knowledgebase, not relying on genomic information. Finally, previous versions of *merlin* (17) used a dedicated tool for this purpose, TRIAGE, which based its annotations on a compilation of manually curated data retrieved from TCDB and predicted transmembrane α-helices. When generated, all reactions were integrated into GSM models, allowing the user to insert any missing information manually.

Other tools are designed to simply address the issue of transporters functional annotation, such as RAST. However, the development of transport proteins’ substrate prediction tools increased in recent years, with tools like The Transporter Substrate Specificity Prediction Server (TrSSP) (18) and the TranCEP (19), that use Support Vector Machine (SVM) models and UniProtKB/Swiss-Prot (20) data in its predictions. The Transporters via Annotation Transfer by Homology (TransATH) provides a different approach (21). This software uses an automatic version of TCDB’s protocol in its annotations. A generic representation of the substrates that each system should transport was obtained through manually pre-processing substrates available in TCDB and the groups to which these substrates belong. Furthermore, the team plans to extract the manual annotations present in TRIAGE’s curated database to improve TransATH’s annotations.

## MATERIAL AND METHODS

*TranSyT* is a Java™ software composed of two main modules, developed considering *TRIAGE’s* limitations. The first module is responsible for regularly gathering, processing, and saving information from different sources. Whereas the second, directed to the end-user, uses that information to identify genes encoding transport proteins in the shortest possible time.

### Data retrieval and transport reactions assembly

*TranSyT*, which also uses TCDB as a primary source of information, can regularly scrape TCDB searching for new entries and updates and retrieving information on transporter systems. Simultaneously, *TranSyT* uses MetaCyc (22) and KEGG to complete information regarding the entries found in TCDB. Descriptions of the families and TC numbers are also retrieved, as often the family reactions include several types of transport.

After gathering all information, the compounds described in the family generic transport reaction, are replaced by the substrates described in the transport systems entry. Additionally, *TranSyT* seeks evidence of direction (in/out) and reversibility in the TC number entry, subfamily, and family descriptions to write the reactions’ equations.

Despite providing the name of the compounds participating in the transport reactions and the recently added feature “Substrate Search Tool”, at the time of *TranSyT’s* implementation, TCDB still did not provide cross-references for substrates described on the page of each TC number. The identification of the compounds used in the reaction is of paramount importance, as stated by Thiele and Palsson (23)at step 14 of the protocol for reconstructing GSM models, if the metabolites do not have identifiers of known databases, these cannot be recognised by scientists and other software tools (23). Thus, to avoid impairing the integration of *TranSyT’s* output with other software, the compounds are identified when generating the transport reactions database. Hence, BioSynth (24), an open-source Bioinformatics software developed by Filipe Liu, is used to accomplish this task. *TranSyT* uses automatic text processing methods to cross BioSynth’s data with substrates’ information retrieved from TCDB to find matches. Each metabolite must follow the process described in figure A1 of supplemental file 1 (supplementary data) to avoid conflicts in the matching process and find matching entries without direct hits. When a match is found, the metabolite is removed from the pipeline. Finally, the databases’ identifiers and chemical formulae for ModelSEED, KEGG, MetaCyc, and BiGG are assigned. Reactions retrieved from MetaCyc already contain information regarding compounds’ identifiers, thus being excluded from this process.

Besides compounds, BioSynth identifies the hierarchical ontologies from MetaCyc, which are used to generate reactions for all hierarchical descendants of the metabolites retrieved from TCDB. For instance, if a TC number describes “sugar” as the transported compound, the identification of all sugars available at MetaCyc is performed, and a transport reaction is generated for each one. However, compounds such as the class “lipid” encompass thousands of descendants. When flagged in *TranSyT’s* configurations, the depth at which these compounds’ descendants are retrieved can be regulated.

*TranSyT* also provides a filter to accept substrates common to ontologies of both TC family and TC number to avoid annotating proteins with false positives by crossing the hierarchical descendants of the TC family’s substrates and the substrates of the TC number. When a match is found, the substrate is saved, and a reaction generated. An example of this process’s importance can be shown using TC number 2.A.40.1.3 (Uniprot accession P75892). The compound assigned to the TC number entry was “Pyrimidines”, and the generic TCDB family reaction described the transport of a “Nucleobase”. Overriding the filter, *TranSyT* generates a total of 114 reactions, as it would create a reaction for all 263 metabolites identified by MetaCyc as descendants of “a Pyrimidine”. However, when the filter is applied, this number drops to three reactions. Although the number of descendants of “a Nucleobase” is nine in MetaCyc, both ontologies only have three shared entries: cytosine, uracil, and thymine. A quick validation using UniProt and EcoCyc using records P75892 and G6517, respectively, confirms that this protein is a transporter with a high affinity for uracil and thymine. However, according to UniProt, this protein does not transport cytosine and seems to have a low affinity to transport xanthine (a purine), a compound not assigned by TCDB. Cytosine appears to be a false positive because it matched the ontologies filter. Nevertheless, the reaction describing xanthine transport can be retrieved from MetaCyc, overcoming TCDB’s misannotation.

During the reactions’ generation process, *TranSyT* seeks evidence of reliability in TCDB’s data, such as descriptions of transport type in the protein’s description and presence of generic TC family reactions. If these criteria are met, MetaCyc’s data is used when in agreement with TCDB, as several reactions from MetaCyc are automatically assigned without human validation. Otherwise, transport reactions for the compounds retrieved from TCDB are generated according to MetaCyc.

All reactions are tested for mass balance, and only balanced transport reactions are accepted as correct. However, in known cases, the reactions require balance correction due to the origin of the annotation. For instance, ATP-binding cassette (ABC) transporters with metabolites formulae retrieved from MetaCyc, BiGG, and ModelSEED will lack a proton. Alternatively, when using KEGG data, the reactions will require a molecule of water. Identified cases are automatically corrected when the reactions are generated. *TranSyT* generates reactions based on evidence of the TC family reaction equation and the description of the TC number, subfamily, family, and superfamily. When no evidence is found, the most straightforward mechanism is assumed (reversible uniport). If symport or antiport evidence is found, but no default reaction is available, a co-transport reaction of the metabolites described in the system together with a proton is generated.

After completing the reactions’ generation and respective mass balance validation, all approved reactions are assigned with a persistent identifier generated by *TranSyT*. The reaction identifiers were designed to be intuitive, i.e., the reaction equation’s content is associated with the identifier, whose structure follows a pattern thoroughly described on page 2 in supplemental file 1. Not all reactions can be intuitively described by the identifier, like a symport of more than two compounds. In such cases, a sequential number is assigned. At this point, cross-references to ModelSEED reactions’ identifiers are also sought. *TranSyT’s* unique identifiers are currently registered under the central registry for life science data Identifiers.org (https://registry.identifiers.org/registry/transyt).

When this process is complete, all information regarding the generated reactions is stored in *TranSyT’s* Neo4j graph database. During the upload process, the organism’s taxonomic identifier is retrieved from the accession number, using NCBI and UniProt APIs, due to the relevance of the phylogenetic information inherent to each transporter system.

### Identification of genes encoding transport systems

*TranSyT’s* second module uses a genome and the respective taxonomy identifier as mandatory input to start the identification process. The taxonomy identifier allows comparing the taxonomy of the organism encoding the reference proteins with the organism submitted by the user. According to Barghash and Helms, 2013 (25), it is more accurate to classify membrane transporters according to TC families than substrates families. Also, according to the same study, two membrane transporters with an expected value score below a threshold of 1E^-8^ on BLAST are likely to share the same TC family, though a threshold of 1E^-4^ could also be considered with caution. Hence, *TranSyT* uses BLAST to identify genes encoding transporter systems, implemented using BLAST+ (25) command-line software available for download at ftp://ftp.ncbi.nlm.nih.gov/blast/executables/blast+/LATEST/, currently in version 2.10. Therefore, all genome sequences are aligned against all sequences present in TCDB’s last retrieved public FASTA file. Additionally, amino acid sequences of proteins involved in phosphotransferase system (PTS) reactions are added to this file to improve the construction of such reactions’ Gene-Protein-Reaction (GPR) associations, such as KEGG KOs K23993 and K02784. *TranSyT’s* default run configurations can be found in Table A1 in supplemental file 1.

After completing the alignments, the next step is to determine which TC family should be assigned to each reference gene by calculating family scores. Such score is defined on *equation 2* in Dias et al. (2017) (7) by constructing a rationale that considers the frequency of hits related to a TC family with the similarity score of such hits. After assigning the family with the highest score to each reference gene, the following step is the association of transport reactions to the genes identified as encoding transport proteins. *TranSyT* uses two different methods to accomplish this task.

The first method accepts all reactions that fulfil the following conditions:

- reactions associated with a TCDB entry must belong to the annotated TC family;
- a TCDB entry hit must have an expected value inferior or equal to the automatic acceptance threshold (0 by default) or belong to the top cluster (10% by default) of the best alignments with an expected value above the acceptance threshold and a given lower threshold (1e-50 by default, which is extremely conservative according to Barghash and Helms, 2013 (25)).

Nevertheless, this method does not consider the hits’ taxonomy due to the evidence of high similarity with the query sequence.

The second method follows an approach similar to *TRIAGE’s* main algorithm for assigning reactions to transport candidate genes. The goal of this method is to find reactions not found by the first method. Hence, this algorithm searches high-frequency reactions among all BLAST hits taxonomically related to the study organism. Reactions with a score above the defined threshold (0.75 by default) are associated with the protein-encoding gene. Detailed descriptions of these calculations and *TranSyT’s* configurations default values can be found on pages 4 and 5 in supplemental file 1.

One of the main features of *TranSyT* is the GPR associations’ algorithm. This software can even search protein complexes formed by multiple subunits. However, the process to find subunits encoded by different genes is not straightforward. *TranSyT* takes advantage of information already available in TCDB regarding protein complexes and the BLAST results to perform this task. Thus, the first step is finding the genes associated with each complex’s subunit, along with the respective bit score. An example of the process is described on page 6 in supplemental file 1.

Using this data, the association is direct in cases where the TC number is related to an isoenzyme or a promiscuous enzyme. The reaction will be encoded by only one protein, encoded by one gene. The algorithm will assign the hit with the highest bit score to each subunit for an enzyme complex. The expected value is used as a tie-breaker for multiple hits with the same bit score. When a match is found, the assigned gene will be removed from other subunits’ results. If the available number of genes is lower than the number of subunits during the process, the associated gene is removed from the GPR, and the next highest scoring gene is processed. This methodology is recursively applied until all subunits are associated with a different gene. If no solution is found, the reaction is removed, as there is no evidence of all subunits in the genome. The results are exported to the Systems Biology Markup Language (SMBL) standard format.

All variables present in *TranSyT’s* calculations are parameterisable through configuration files, and the user can override input settings. When the process is finished, the results provide a simple annotation of each gene’s proteins and reactions. Otherwise, when integrating the output with a GSM model, *TranSyT* can use the list of compounds present in the model to filter the reactions that would not have flux.

## RESULTS AND DISCUSSION

Three different case studies assessed *TranSyT’s* performance. The first case study uses Biolog data from 103 different genomes to analyse with experimental data, whether *TranSyT* can assign transport reactions related to Biolog’s report compounds. It also evaluates which set of parameters best fit the results to be used in the test cases. The second case study analyses RAST’s annotations for 1671 genomes, comparing, when possible, the type of transport and a sample of compounds between both annotations. The third case study thoroughly compares the transport reactions present in the iML1515 (26) model of *Escherichia coli str. K-12 substr. MG1655* (*E. coli*) with *TranSyT’s* annotations for the same genome.

Despite providing support to 4 different databases, ModelSEED is used as the cornerstone of the internal database as the software is integrated in KBase. Thus, the results were generated using ModelSEED identifiers.

### Growth phenotype data

Biolog provides experimental data regarding the substrates used by an organism, indicating that the organism ought to be endowed with mechanisms that transport such compounds across the external membrane. The report is composed of 103 different organisms with growth/no-growth indications for 67 (64 + 3 particular cases) different substrates. The rationale for using these particular cases is detailed in sheet B3 of supplemental file 2.

*TranSyT* was executed using different sets of parameters for both methods of identification (ran independently). Method one was implemented using default parameters, only changing the percentage (*p*) of best BLAST results to accept: 0%, 5%, 10% and 20%. The second identification method also used default parameters, except for the *α* value set to 0.5 and 0.75.

Table 1 contains a summary of the average results for each method independently and merged with the other approach for each set of parameters. The combined results present a negligible increase in the assignment of reactions to growth compounds when M_1*p*_ = 0% and M_2*α*_ = 0.5. However, when increasing M_2*α*_ to 0.75, the rise is greater than 7% when M_1*p*_ = 0%, though less than 1% when M_1*p*_ > 0%. A careful analysis of the results shows that as expected, the largest rise occurs for genomes with smaller representation in TCDB. The most extreme case is *Marinobacter adhaerens* HP15, with an increase of 20%. This organism has one entry in TCDB, and its genus is not well represented, with 15 entries only. When M_1*p*_ > 0% and M_2*α*_ = 0.5, the same increase is not verified, with very few organisms having an increase in the assignment of reactions to growth compounds.

**Table 1.**
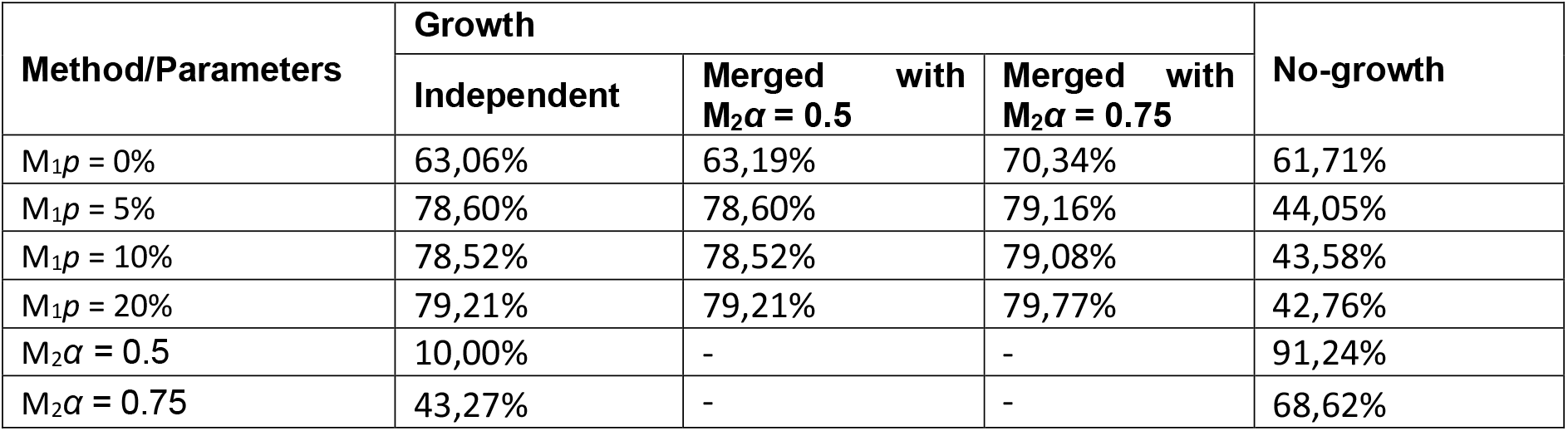
Average growth/no-growth compounds found. Growth substrates contain at least one reaction. No-growth substrates had no reactions assigned.

| Method/Parameters | Growth |  |  | No-growth |
| --- | --- | --- | --- | --- |
| | Independent | Merged with $M_2\alpha = 0.5$ | Merged with $M_2\alpha = 0.75$ | |
| $M_1p = 0\%$ | 63,06% | 63,19% | 70,34% | 61,71% |
| $M_1p = 5\%$ | 78,60% | 78,60% | 79,16% | 44,05% |
| $M_1p = 10\%$ | 78,52% | 78,52% | 79,08% | 43,58% |
| $M_1p = 20\%$ | 79,21% | 79,21% | 79,77% | 42,76% |
| $M_2\alpha = 0.5$ | 10,00% | - | - | 91,24% |
| $M_2\alpha = 0.75$ | 43,27% | - | - | 68,62% |

As expected, the increase of M_1*p*_ and M_2*α*_ drops the number of True Negatives, i.e. non-growth substrates without transport reactions assigned by *TranSyT*. Although these compounds are not associated with the organism’s growth, it does not indicate that these are not transported across the membrane to participate in other metabolic functions. Thus, M_1*p*_ = 10 and M_2*α*_ = 0.75 were selected for the next test cases, not being over or under conservative.

Five substrates had no reactions assigned: L-Hydroxyproline, Quinic Acid, α-Ketobutyric Acid, b-Methyl-D-Glucoside, and gly-pro-L. As only α-Ketobutyric Acid and Quinic Acid are present in *TranSyT’s* database, the non-growth results might be biased.

Additional information about the results here provided can be found in supplemental file 2.

### Analysis of a large set of phylogenetic diverse genomes

The second test case uses RAST and RefSeq annotations for 1671 genomes. By collecting the annotation for the genes assigned as encoding transport proteins by *TranSyT*, it was possible to compare the compounds in transport and the transport types. Only genes annotated as transporters by *TranSyT* are retrieved as despite other transporters might be annotated by RAST and RefSeq, *TranSyT* generates transport reactions with relevant functions to GSM models. Thus, a dataset of 216890 genes was collected for analysis. By searching keywords associated with transport types in the annotation description, the following categories were assigned to the transport proteins: Simple, ABC, PTS, Cofactor, and oxidation-reduction (Redox). Simple is the most generic category, as it encompasses the simplest transport types: Uniport, Symport, and Antiport. These three were grouped in the same category as it was challenging to distinguish transporters characterised as “Major Facilitator Superfamily” or “porin”. Nevertheless, it was not possible to infer the transport type for around 27% of RAST’s annotations and 35% of RefSeq’s annotations, as it was not possible to automatically assume a transport type (ex: Shikimate transporter). Moreover, regarding RefSeq’s dataset, almost 10% of the entries were not annotated.

The filtered datasets included 150949 and 134116 transporters for RAST and RefSeq, respectively.

According with Figure 1, all categories match over 92% of RAST and 89% of RefSeq’s annotations. One of the cases analysed was regarding the gene STERM_RS12595, in which RAST’s annotation is ABC transporter. In this case, *TranSyT’s* BLAST results assigned the gene to TC family 9.B.20. Entries assigned in TC class 9 are usually incompletely characterised and in many instances lacking a TC family generic reaction, causing *TranSyT* to flag TCDB’s data as unreliable for such cases and use MetaCyc instead. In this specific case, MetaCyc’s reaction was a Uniport created without human supervision. However, the compound in transport was in agreement with RAST’s annotation. For the same gene, RefSeq’s annotation had no reference to the transport type in its annotation and was therefore not included in the dataset.

**Figure 1.**
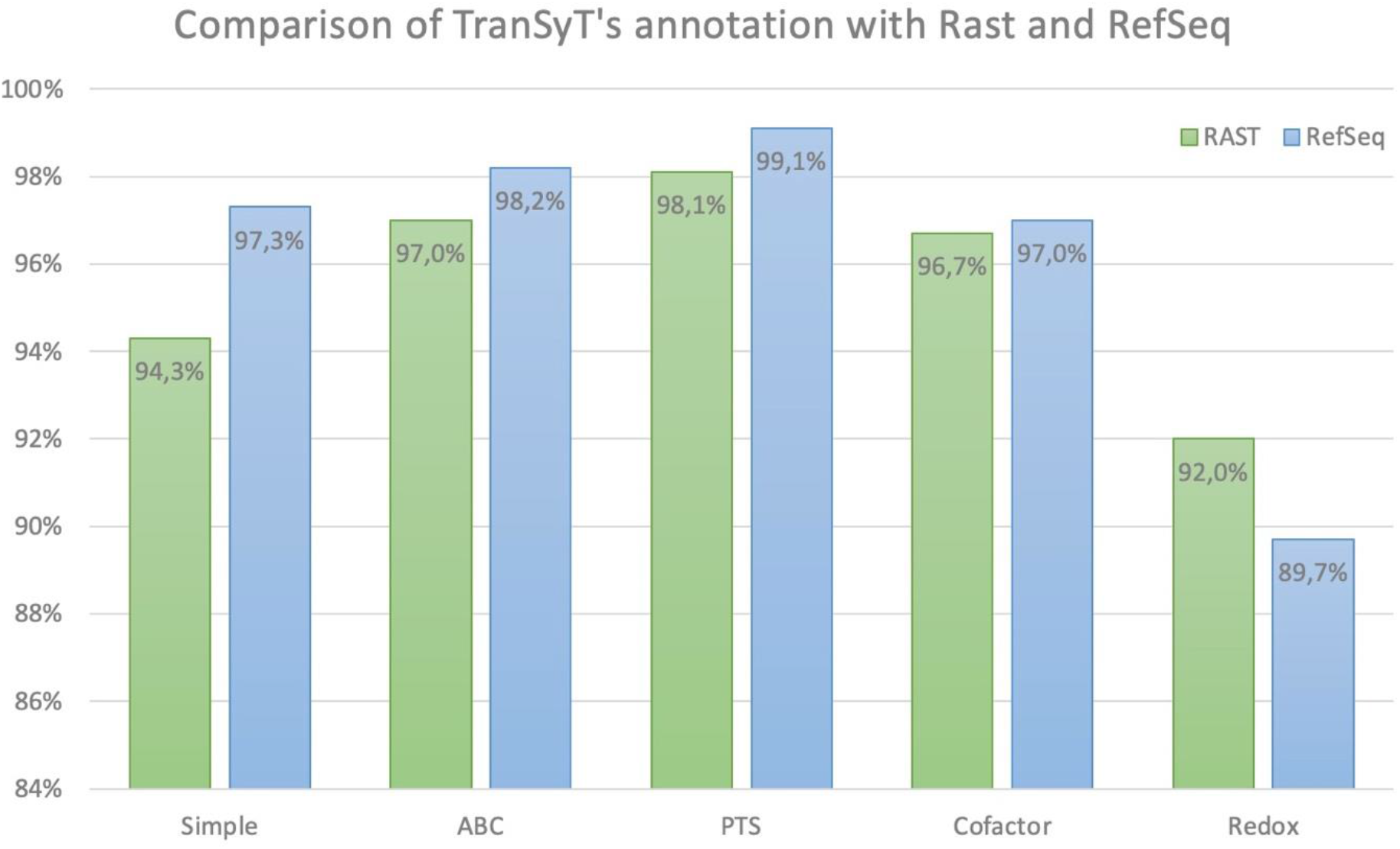
Comparison of TranSyT’s genes transport type annotation against RAST and RefSeq filtered datasets. The bars represent the percentage of genes annotated by TranSyt with identical annotation on either RAST or RefSeq.

A similar approach was used to compare the compounds in the filtered dataset. However, in this case, only RAST annotations were used as the annotations are generally more descriptive, and all genes had an annotation. Cytosine, sulfate, acetate, putrescine, copper, nitrate, and cellulose were selected for the study.

As shown in Figure 2, cytosine, copper, and nitrate had the lowest match rate, whereas the remaining compounds scored above 86% in agreement with RAST’s annotation. A more detailed analysis of *TranSyT’s* annotations for cytosine shows that 24.6% of the cases annotated as transporting this compound had the description “Cytosine/purine/uracil/thiamine/allantoin permease family protein” assigned by RAST and TC number 2.A.39.3.4 by *TranSyT*. For this family, TCDB only describes allantoin as transporting compound. MetaCyc’s reaction is also in agreement with TCDB, describing the homolog as “allantoin permease”.

**Figure 2.**
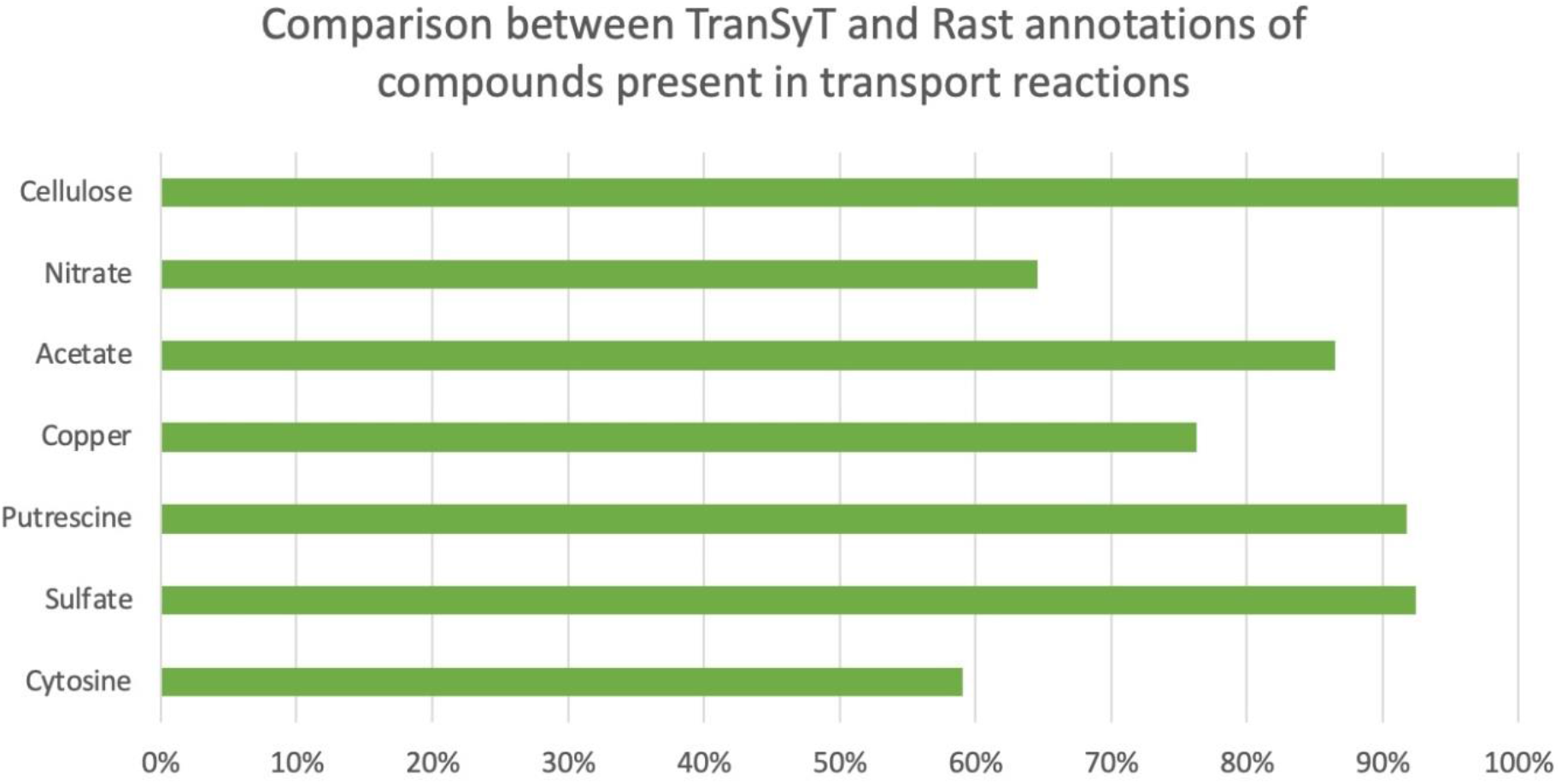
Comparison of genes associated with the transport of 7 randomly selected metabolites between TranSyT and RAST. These metabolites were selected from the shared pool of metabolites obtained with the previous assessments. The bars represent the percentage of genes annotated by TranSyT with identical annotation on RefSeq for each compound.

A second case was analysed for cytosine in which RAST had the same annotation as before. However, *TranSyT* assigned the TC number 2.A.39.3.14, for adenine, guanine, uracil, and allantoin transport, though in this case, MetaCyc provided no information. Nevertheless, as TCDB provides cross-references to literature sources, it was possible to find in Schein’s (2013) (27) that this protein cannot transport cytosine and uridine. This situation was observed in 6.6% of genes.

Regarding copper annotations, in 15% of the cases, RAST assigned the genes as “Copper-translocating P-type ATPase (EC 3.6.3.4); Lead, cadmium, zinc, and mercury transporting ATPase (EC 3.6.3.3) (EC 3.6.3.5)”. With the advent of EC number class 7, the aforementioned EC numbers were transferred to EC 7.2.2.9, EC 7.2.2.21, and EC 7.2.2.12, respectively. Running a BLAST against such genes and combining the results with UniProt data, it was possible to determine that the TC numbers with the best results in the alignment are associated with EC numbers 7.2.2.21 and 7.2.2.12. These TC numbers do not contain copper in their descriptions. A relaxation of the parameters would eventually allow *TranSyT* to include TC numbers with lower similarity related to EC number 7.2.2.9, and consequently, copper reactions.

A third case was analysed for nitrate, where it was found that for 10% of the entries RAST annotated these genes as “Nitrate/nitrite transporter”. When running a BLAST of the same genes on NCBI, it was possible to confirm *TranSyT’s* annotations of “putative tartrate transporter” genes.

### Comparison with published model

*TranSyT’s* annotations for *E. coli* str. K-12 substr. MG1655 were assessed against the iML1515 GSM model. This model contains 1516 genes, 2712 reactions, 1877 metabolites, and uses BiGG identifiers for its reactions and compounds. However, for consistency with the previous studies, ModelSEED was used in TranSyT’s annotations.

Transport reactions information per gene was retrieved from the model using COBRA toolbox (28). All reactions with metabolites in more than one compartment were considered transporters. Therefore, the model contains 833 transport reactions, 780 of which distributed by 499 genes, with 53 transport reactions without a valid GPR.

To understand the importance of giving the set of compounds present in the model as input to *TranSyT*, both scenarios were assessed. The first test case was generated without using the metabolites filter, resulting in *TranSyT* annotating 670 genes as encoding transport proteins and 10164 transport reactions. Of the genes associated with transporters in the model, *TranSyT* did not include results for 69 genes, though identifying 240 new genes not included in the model or assigned with non-transport functions.

The second method was performed using the filter, creating 1626 transport reactions, 84% less than the former methodology. The number of genes with transport reactions was significantly lower (572); unexpectedly, only two more genes (71 in total) were annotated as transporters in iML1515 but not by TranSyT. A detailed analysis of the results obtained is available in Figure 3.

**Figure 3.**
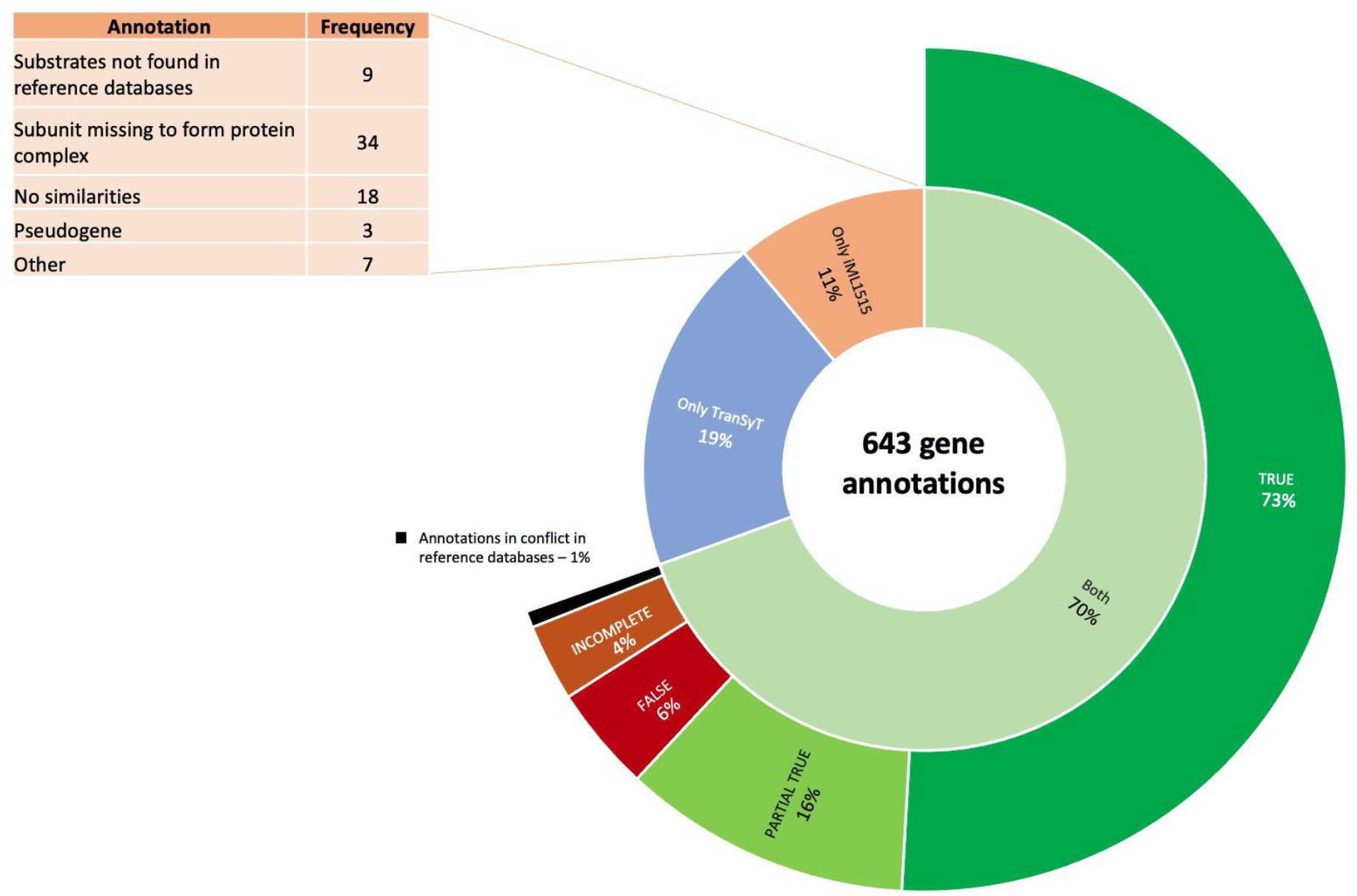
TranSyT *results for* E. coli *str. K-12 substr. MG1655 using only compounds present in the iML1515 model.The figure represents the percentage of gene annotations present only in the model, in TranSyT, and both of them, as well as the comparison of the common transport reactions*.

*TranSyT* and iML1515 share a set of genes (447) for which both have transport reactions. Here, 73% match perfectly the transport reactions. A partial match was obtained for 16%, in which *TranSyT* has reactions for a significant portion of the compounds transported in the model. There is usually no clear evidence in the reference databases that the model’s transport reactions should be associated with such genes. Regarding the 4% gene annotations assessed as incomplete, the compounds are mentioned in the reference databases, but not described in TCDB’s substrates and MetaCyc. Whereas 6% of the gene annotations are utterly different in *iML1515 and TranSyT*. In these situations, *TranSyT* is usually supported by the evidence found in the reference databases. Finally, there are misannotations both in TCDB or MetaCyc that lead to creating incorrect transport reactions. Such genes represented 1% of the annotations and were assessed as in “conflict”, where the references have contradictory annotations.

*TranSyT* assigned reactions to 125 genes, not available in iML1515. A sample (33 genes) was manually curated, revealing that the reference databases support *TranSyT’s* classification. The complete annotations are available in sheet C2 of supplemental file 3.

A critical feature in *TranSyT* is the assignment of GPR rules to protein complexes. An example is iML1515’s reaction R_NADH16pp, in which the GPR includes 13 subunits. The same result was obtained in *TranSyT*, which annotated the reaction with TC number 3.D.1.1.1 and the same set of genes. A second example is the PTS reaction R_MANptspp, associated with five subunits in the model and three subunits in TCDB (TC number 4.A.6.1.1). In such cases, *TranSyT* uses KEGG to find the missing subunits. In this example, KEGG KOs K23993 and K02784, related to TC families 8.A.7 (The Phosphotransferase System Enzyme I Family) and 8.A.8 (The Phosphotransferase System HPr Family), respectively. Additionally, *TranSyT* also found a second alternative for the same reaction for TC number 4.A.1.1.14, with three subunits.

Finally, the model analysis showed that the iML1515 model does not provide gene rules for 53 transport reactions. *TranSyT* matched six of these reactions perfectly and generated gene rules for each one. A more thorough analysis showed that *TranSyT* matched partially 12 of such reactions, in which the difference is either the transport type or reversibility. The remaining 35 reactions were not available in *TranSyT’s* output. Detailed information about these reactions is available in sheet C5 of the supplemental file C.

## CONCLUSIONS

A combination of all case-studies’ results allows understanding that *TranSyT’s* only limitation is the restricted number of different organisms available in TCDB, having the highest performance for organisms taxonomically close to *E. coli* and *Homo sapiens*, for which TCDB is biased. Hence, *TranSyT* uses the second classification method to overcome this issue, which should never replace the primary classifier, instead as a counterpart. Nevertheless, additional highly curated sources of transport data should be integrated in the future. Also, tools such as TrSSP and TranCEP could be integrated into *TranSyT* as these approaches use other algorithms and databases, extending *TranSyT’s* coverage and improving performance.

Without the metabolites filter, TranSyT returns an excessive amount of possible transport reactions. Nevertheless, this can be seen as a useful resource in cases where it might be interesting to investigate an organism’s capability to transport a specific metabolite across its membranes. Since *TranSyT’s* reactions already provide information regarding the direction of the reactions, the integration of membrane localisation information can also be easily achieved using third-party software.

*TranSyT* is currently available as a web service at https://transyt.bio.di.uminho.pt/, a plugin in *merlin* v4 (29), and a beta app in KBase (30).

## AVAILABILITY

*TranSyT’s* web service - https://transyt.bio.di.uminho.pt/

*merlin* v4 - https://merlin-sysbio.org/

KBase - https://www.kbase.us/

GitHub – https://github.com/BioSystemsUM/transyt

## SUPPLEMENTARY DATA

Supplementary Data are available at NAR online.

## ACKNOWLEDGEMENT

The submitted manuscript has been created by UChicago Argonne, LLC as Operator of Argonne National Laboratory (‘Argonne’) under Contract No. DE-AC02-06CH11357 with the U.S. Department of Energy. The U.S. Government retains for itself, and others acting on its behalf, a paid-up, nonexclusive, irrevocable worldwide license in said article to reproduce, prepare derivative works, distribute copies to the public, and perform publicly and display publicly, by or on behalf of the Government. The Department of Energy will provide public access to these results of federally sponsored research in accordance with the DOE Public Access Plan.The authors would like to acknowledge project 22231/01/SAICT/2016: “Biodata.pt – Infraestrutura Portuguesa de Dados Biológicos”, supported by Lisboa Portugal Regional Operational Programme (Lisboa2020), under the PORTUGAL 2020 Partnership Agreement, through the European Regional Development Fund (ERDF). The authors would also like to acknowledge the Portuguese Foundation for Science and Technology (FCT) for providing a PhD scholarship to E. Cunha (DFA/BD/8076/2020). Oscar Dias acknowledges FCT for the Assistant Research contract obtained under CEEC Individual 2018.

