## Supplemental file 1 for "*TranSyT*, an innovative framework for identifying transport systems"

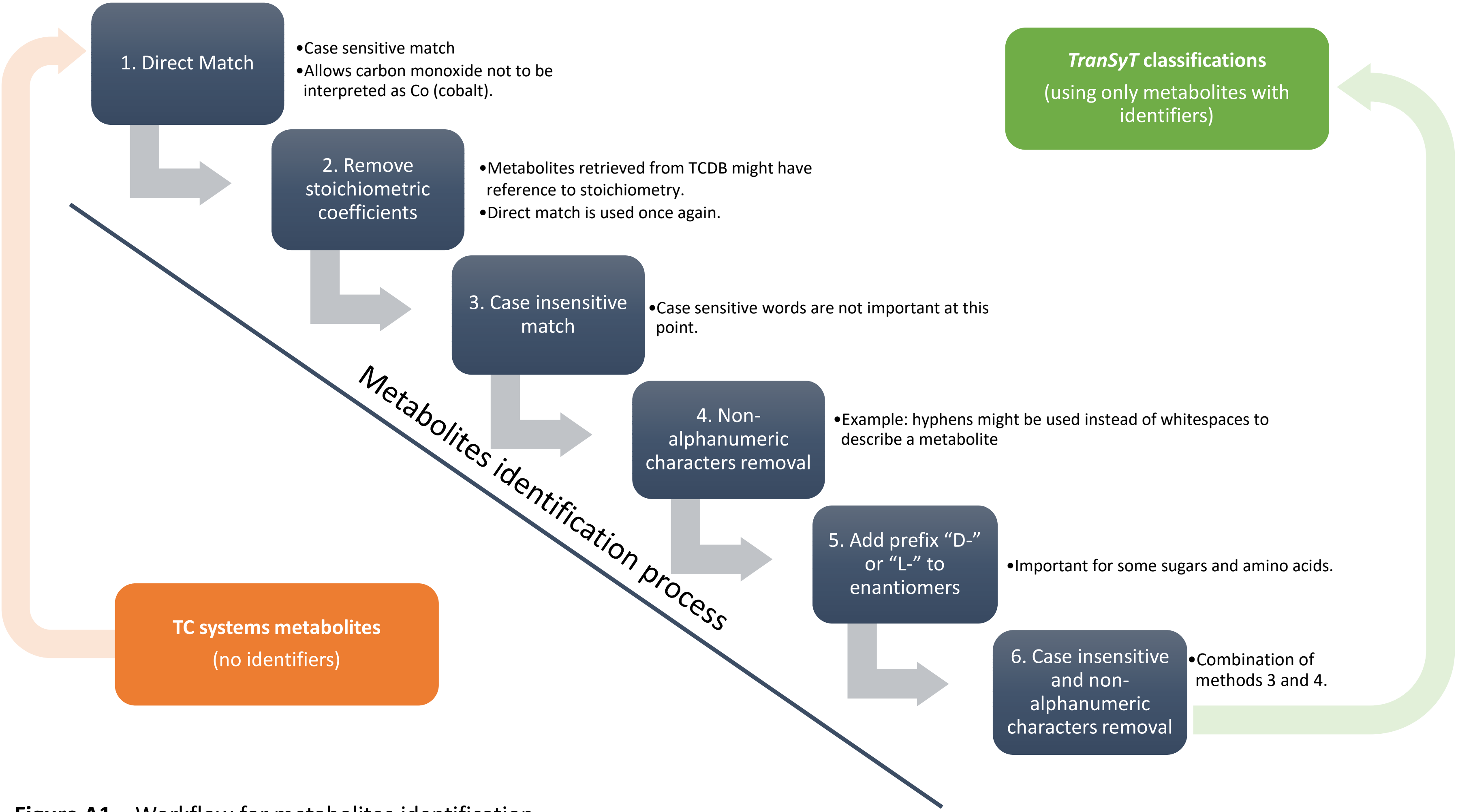


**Figure A1 –** Workflow for metabolites identification

### Identifiers generator – Example: TR0000067

**T [A-Z] \d \d \d{5}**

**Always a "T" in the beginning** ← T

[A-Z] → **Reversibility and direction**

| | |
|---|---|
| Reversible | R |
| In to out | I |
| Out to in | O |
| Default | Z |

\d → **Transport type**

| | |
|---|---|
| Uniport | 0 |
| Symport | 1 |
| Antiport | 2 |
| ABC | 3 |
| PTS | 4 |
| Cofactor | 5 |
| Redox | 6 |
| Default | 9 |

\d → **Co-transported metabolite***

| | |
|---|---|
| Hydrogen | 0 |
| Sodium | 1 |
| Potassium | 2 |
| Calcium | 3 |
| Magnesium | 4 |
| Chloride | 5 |
| Zinc | 6 |
| Iron | 7 |
| Default | 9 |

*only if symport or antiport with 2 metabolites

\d{5} →

- **If transport type is uniport, symport(2 metabolites), antiport (2 metabolites):**
  - modelSEED identifier of the metabolite in transport.
- **Else:**
  - persistent sequential number is generated.

NOTE: *TranSyT*'s unique identifiers are currently registered under the central registry for life science data Identifiers.org.
https://registry.identifiers.org/registry/transyt

**Table A1 –** *TranSyT*'s default configurations

| | | |
|---|---|---|
| ❑ **Alignments parameters*** | | |
| E-value threshold | $1e^{-20}$ | BLAST E-value threshold. |
| Minimum query coverage | 80% | BLAST query coverage threshold. |
| Similarity score | 30% | Parameter used to filter BLAST results |
| Bit score | 50 | BLAST bit score threshold. |
| Scoring matrix | BLOSUM62 | Substitution matrix used by BLAST. |
| ❑ **Reactions annotation method-1** | | |
| E-value auto-accept threshold | 0 | The value below which all hits are accepted as possible transporters. |
| % of top results acceptance | 10% | Portion of best hits that should be accepted for each query gene. |
| E-value threshold | $1e^{-50}$ | E-value above which all results should be disregarded by method-1. |
| ❑ **Reactions annotation method-2** | | |
| α-value | 0.75 | α-value of reactions annotation equation. Used to balance the weight between frequency and taxonomy scores. |
| β-value | 0.3 | β-value of taxonomy score equation. Used as penalty for reactions that are associated to a low number of homologous genes. |
| Minimum hits penalty | 2 | Minimum hits to avoid low-hits penalty in taxonomy score equation. |
| Score threshold | 0.75 | Score above which a reaction is accepted. |
| ❑ **Other** | | |
| TC families α-value | 0.4 | Parameter used to balance the weight between frequency and similarity scores during TC family calculations. |

*The default parameters for BLAST were set considering the Pearson's study in An Introduction to Sequence Similarity ("Homology") Searching. This study indicates that bit scores of 40 and E-values ≤ 1E-3 are significant for databases with less than 7 000 entries. Since the E-value is dependent on the size of the database, an increase of 10 to the bit score increases the significance in a factor of $2^{10}$, thus a bit score of 50 would be significant for a database with less than 7 million entries (TCDB's FASTA file currently contains almost 19000 entries). Hence, a low E-value, a high bit score and 80% coverage of the query sequence when querying a database as small as TCDB, can be regarded as a conservative approach. This strategy will increase the number of false negatives rather than the number of false positives. Nevertheless, all values are configurable.

#### Family score calculations

The family score is calculated using Equation 1, that uses the α value to balance the weight of the frequency ($Score_{Freq}$) of the families in the BLAST results and the respective similarities ($Score_{Sim}$). α can be set with values ranging from 0 to 1, being the default 0.4 which slightly favors the similarity without disregarding the frequency.

$$Score = \alpha \times Score_{Freq} + (1 - \alpha) \times Score_{Sim} \quad \Rightarrow \quad Equation\ 1$$

The frequency score is calculated by dividing the number of hits of a specific TC family *F*, by the total number of BLAST hits. Thus, $F_i$ refers to the $i^{th}$ BLAST record, and *H* the total number of hits.

$$Score_{Freq} = \frac{\sum_{i=1}^{H} F_i \times Vf_i}{\sum_{i=1}^{H} F_i} \quad \text{where,} \quad Vf_i = \begin{cases} 1, & if\ record\ i\ belongs\ to\ family \\ 0, & otherwise \end{cases} \quad \Rightarrow \quad Equation\ 2$$

Similarly, the similarity score is calculated by dividing the sum of the similarities of the hits belonging to family *F* by the sum of the similarities of all hits. $S_i$ refers to the similarity of the $i^{th}$ BLAST record, and *H* the total number of hits.

$$Score_{Sim} = \frac{\sum_{i=1}^{H} S_i \times Vfs_i}{\sum_{i=1}^{H} S_i} \quad \text{where,} \quad Vfs_i = \begin{cases} 1, & if\ record\ i\ belongs\ to\ family \\ 0, & otherwise \end{cases} \quad \Rightarrow \quad Equation\ 3$$

#### Reactions score calculations

The reaction score in calculated with a similar equation used for the family score. In this case, for each gene *g*, the frequency of a reaction *r* within the homologous protein records is used as frequency score ($Score_{Freq}$), while the taxonomy score ($Score_{Sim}$) calculation is based on the common taxonomy between query and homologous hits. Similarly, both scores are weightned by the parameter α that can be set with values ranging from 0 to 1.
Since TCDB contains mostly records belonging to Bacteria, 0.75 is used as default value, however, if the case study organism does not belong to this domain, it is recommended the reduction of this value.

$$Score = \alpha \times Score_{Freq} + (1-\alpha) \times Score_{Tax} \quad \Rightarrow \quad Equation\ 4$$

The frequency score is calculated by dividing the similarity of all homologous proteins promoting the reaction by the sum of the similarities of all BLAST hits, for each protein encoding gene. Thus, $S_i$ refers to the similarity of the $i^{th}$ BLAST hit encoding the reaction being scored, and *H* the total number of hits.

$$Score_{Freq} = \frac{\sum_{i=1}^{H} S_i \times Vr_i}{\sum_{i=1}^{H} S_i} \quad \text{where,} \quad Vr_i = \begin{cases} 1, & if\ reaction\ r\ is\ represented\ in\ record\ i \\ 0, & otherwise \end{cases} \quad \Rightarrow \quad Equation\ 5$$

The taxonomy score is calculated by dividing the taxonomy frequency $t_i$ by the maximum taxonomy $M_T$(number of taxa of the organism in study), multiplied by the frequency of the protein hits promoting the reaction and a penalty. This penalty is applied reactions that are associated to a low number of homologous genes by multiplying $p_r$ with β (this value ranges from 0 to 1 and is set by default as 0.3). The taxonomy frequency is calculated by counting the common taxa between the case study and the organism to which the $i^{th}$ homologous record belongs.

$$Score_{Tax} = \frac{\sum_{i=1}^{H} t_i \times Vr_i \times (1 - p_r \times \beta)}{M_T \times \sum_{i=1}^{H} Vr_i} \quad \text{where,} \quad p_r = \begin{cases} 0, & if\ \sum_{i=1}^{H} Vr_i \geq Min_{Hits} \\ Min_{Hits} - \sum_{i=1}^{H} Vr_i, & otherwise \end{cases} \quad \Rightarrow \quad Equation\ 6$$

#### *TranSyT*'s method of assignment of Gene-Protein-Reaction (GPR) rules to transport reactions in case of protein complex with multiple hits.

Table A2 contains the results of the alignment of the TCDB records against the query genome.

TranSyT's database contains the information that the protein assigned to TC number Y is composed of 3 different subunits: A, B and C.

A, B, and C have similarity with the query genes represented in the table A2.

TranSyT uses the information from the table and verifies for each accession, which of the query genes has most similarity, trying to achieve an optimal solution. In case an accession has the same similarity score for two query genes, the lowest E-value is used as tie-breaker. Similarly, two different accessions cannot be assigned to the same query gene.

In case a solution is not possible, this means that one of the subunits is missing, and therefore the protein is not completed. The reaction is excluded.

From the example table of reaction X, the output of this rationale is: A = Gene_1; B = Gene_3; C = Gene_2
GPR rule = Gene_1 and Gene_2 and Gene_3

In cases where the TC number does not have multiple subunits associated, the direct assignment of the best similarity score with E-value tie-breaker is used.
Example: If TC number Y had only accession A associated with the same 4 query genes, GPR would be Gene_1

**Table A2 –** Example containing similarity results for reaction X.

| TC Number | Accession | Query Gene | Similarity |
|---|---|---|---|
| Y | A | Gene_1 | 0,7 |
| | | Gene_2 | 0,5 |
| | | Gene_3 | 0,66 |
| | | Gene_4 | 0,58 |
| | B | Gene_1 | 0,25 |
| | | Gene_3 | 0,8 |
| | C | Gene_2 | 0,5 |
| | | Gene_3 | 0,25 |
